# IUPAC Consensus References Improve Short-Read Variant Detection in Clinically Challenging Regions: A Stratified Benchmarking Study with BurdenBench

**DOI:** 10.64898/2026.08.12.744558

**Authors:** Akzam Saidin, Michael G. Ricos, Leanne M. Dibbens

**Affiliations:** Epilepsy Research Group, School of Pharmacy and Biomedical Sciences, College of Health, Adelaide University, South Australia 5000, Australia; Novocraft Technologies, Petaling Jaya, Malaysia

**Keywords:** reference bias, consensus reference, IUPAC degenerate codes, GRCh38, variant calling, benchmarking, clinical genomics, pangenome, whole-genome sequencing

## Abstract

**Motivation:** Reference bias depresses variant detection in low-mappability regions, segmental duplications and the major histocompatibility complex (MHC) — precisely the regions of greatest clinical relevance. Existing benchmarks rely on aggregate precision, recall and F1 metrics that obscure the absolute true-positive and false-positive counts that determine laboratory workload. No study has systematically evaluated IUPAC consensus references for short-read whole-genome sequencing (WGS) variant calling across Genome in a Bottle (GIAB) stratifications, multiple allele-frequency thresholds and multiple variant callers.

**Results:** We aligned 30× WGS from three GIAB samples to IUPAC consensus references (allele frequency ≥10% and ≥30%) using the ambiguity-aware aligner novoAlign, benchmarking against BWA-MEM/GRCh38 and novoAlign/GRCh38 baselines across BCFtools, FreeBayes and GATK HaplotypeCaller. SNV recall increased by 3.1–3.9 percentage points (pp) in low-mappability regions and 1.8–3.1 pp in segmental duplications; INDEL recall rose by 4.5–5.8 pp and 2.4–3.8 pp, respectively, with similar gains in the MHC and challenging medically relevant genes (CMRG). Decomposition analysis showed that the aligner change drove most INDEL gains, while IUPAC encoding contributed additional SNV-specific improvement. We introduce BurdenBench, an open-source framework that computes net benefit and region-size-normalised metrics directly from standard hap.py outputs, revealing divergent caller-specific trade-off profiles that are invisible to aggregate F1: FreeBayes showed the most favourable precision–recall balance in low-mappability regions, while GATK achieved positive net benefit in the MHC. A controlled comparison using an identical variant set showed severe recall and precision losses for SALT (a published SNP-aware dual-index aligner) across all three callers, supporting the value of preserving linear reference structure. Pan-human and population-specific consensuses performed within 0.2 pp of one another. All findings are descriptive and hypothesis-generating from three samples.

**Availability and implementation:** To mitigate potential bias associated with software developed by an author’s employer, primary hap.py outputs and derived burden metrics were independently verified by co-authors with no affiliation to that employer. BurdenBench (v1.0.0) is implemented in Python (pandas, numpy; Python ≥3.7) and freely available under the MIT licence at https://github.com/akzam/BurdenBench, including raw hap.py outputs and an audit trail enabling independent recomputation without a novoAlign licence. novoAlign and novoUtil (version 4, Novocraft Technologies) are commercial software with no-cost academic trial licences.

**Supplementary information:** Supplementary tables, figures and methods are available online.

## 1 Introduction

The first human genome draft established a universal coordinate system underpinning modern clinical genomics (Lander et al., 2001), but the limitations of a single, largely haploid linear reference have become increasingly consequential as sequencing enters routine precision medicine (Ballouz et al., 2019; Majidian et al., 2023). A central technical driver is reference bias: short-read aligners favour the reference allele, producing misplaced alignments and systematic errors in small-variant detection, particularly in polymorphic and repetitive loci (Lin et al., 2024; Oliva et al., 2021). Clinically, this can depress heterozygous genotype quality and reduce sensitivity for pathogenic variants, with patients whose genomes diverge most from the reference disproportionately affected (Lin et al., 2024; Gunther and Nettelblad, 2019; Ballouz et al., 2019). Assembly errors in GRCh38 compound the problem, and performance benchmarked in high-confidence regions may overestimate accuracy in the clinically relevant loci where bias is most pronounced (Dwarshuis et al., 2024; Behera et al., 2023).

Emerging reference strategies promise better representation of human diversity. T2T-CHM13 resolves gaps and misassemblies in GRCh38, while the HPRC draft pangenome captures haplotypic diversity absent from any single linear reference (Nurk et al., 2022; Liao et al., 2023). Clinical translation, however, has been slow: reanalysing legacy cohorts is costly, CLIA/CAP validation burdens are substantial, and graph-based models face interoperability gaps and tooling immaturity (Majidian et al., 2023; Lovell and Grimwood, 2022). GA4GH and GIAB stratified benchmarking resources now enable rigorous evaluation in difficult regions, providing the infrastructure needed to quantify gains from reference improvements (Dwarshuis et al., 2024).

A pragmatic, pipeline-compatible mitigation is the consensus reference: common alleles are substituted into a linear reference to reduce mismatch penalties while preserving coordinate compatibility with existing tools (Ballouz et al., 2019; Chen et al., 2021). IUPAC degenerate codes can encode these common variants without altering contig lengths, and ambiguity-aware aligners such as novoAlign score IUPAC positions as matches, lowering mismatch penalties for heterozygous reads (Johnson, 2010). This approach has been validated in ancient DNA read mapping, where novoAlign’s IUPAC scoring improved alignment fidelity against consensus references for damaged reads (Oliva et al., 2021), and in RNA-seq, where a pan-human consensus reduced mapping errors two- to three-fold for reads overlapping homozygous variants (Kaminow et al., 2022). For clinical genomics, IUPAC consensus alignment is attractive because it preserves GRCh38 linear coordinates, requires no graph-aware tooling, and can be validated incrementallyalongside existing pipelines. novoAlign is currently the only publicly available aligner supporting true IUPAC degenerate base scoring at the reference-index level for standard short-read WGS; DRAGEN’s graph mapper uses a related but architecturally distinct hybrid approach (Illumina, 2021), and mrFAST’s SNP-aware mode is non-functional in current releases (sfu-compbio/mrsfast issue #14).

No study has systematically evaluated IUPAC consensus references for short-read WGS variant calling across GIAB stratifications, multiple allele-frequency thresholds and multiple callers. Furthermore, existing benchmarks rely on aggregate metrics (precision, recall, F1) that do not expose absolute true-positive and false-positive counts: two scenarios with identical F1 gains may differ by orders of magnitude in practical impact (Majidian et al., 2023). Here, we address both gaps. We report a systematic exploratory assessment using three GIAB samples (HG001, HG002, HG005) and three variant callers, evaluating precision–recall trade-offs across GIAB stratifications and decomposing aligner and IUPAC contributions. We compare IUPAC consensus against SALT (a published SNP-aware dual-index aligner) across all three callers using an identical variant set, and we introduce BurdenBench, a reproducible, open-source framework that extends standard hap.py outputs to report absolute true-positive and false-positive counts, net benefit and region-size-normalised metrics. All findings are descriptive and hypothesis-generating; the sample size reflects current GIAB truth-set availability rather than a design choice.

## 2 Materials and methods

### 2.1 Workflow overview

Common SNPs were extracted from population databases and substituted into GRCh38.p13 as IUPAC degenerate codes; reads were aligned with novoAlign, variants called with three callers, and results benchmarked against GIAB truth sets (Fig. 1).

**Fig. 1.**
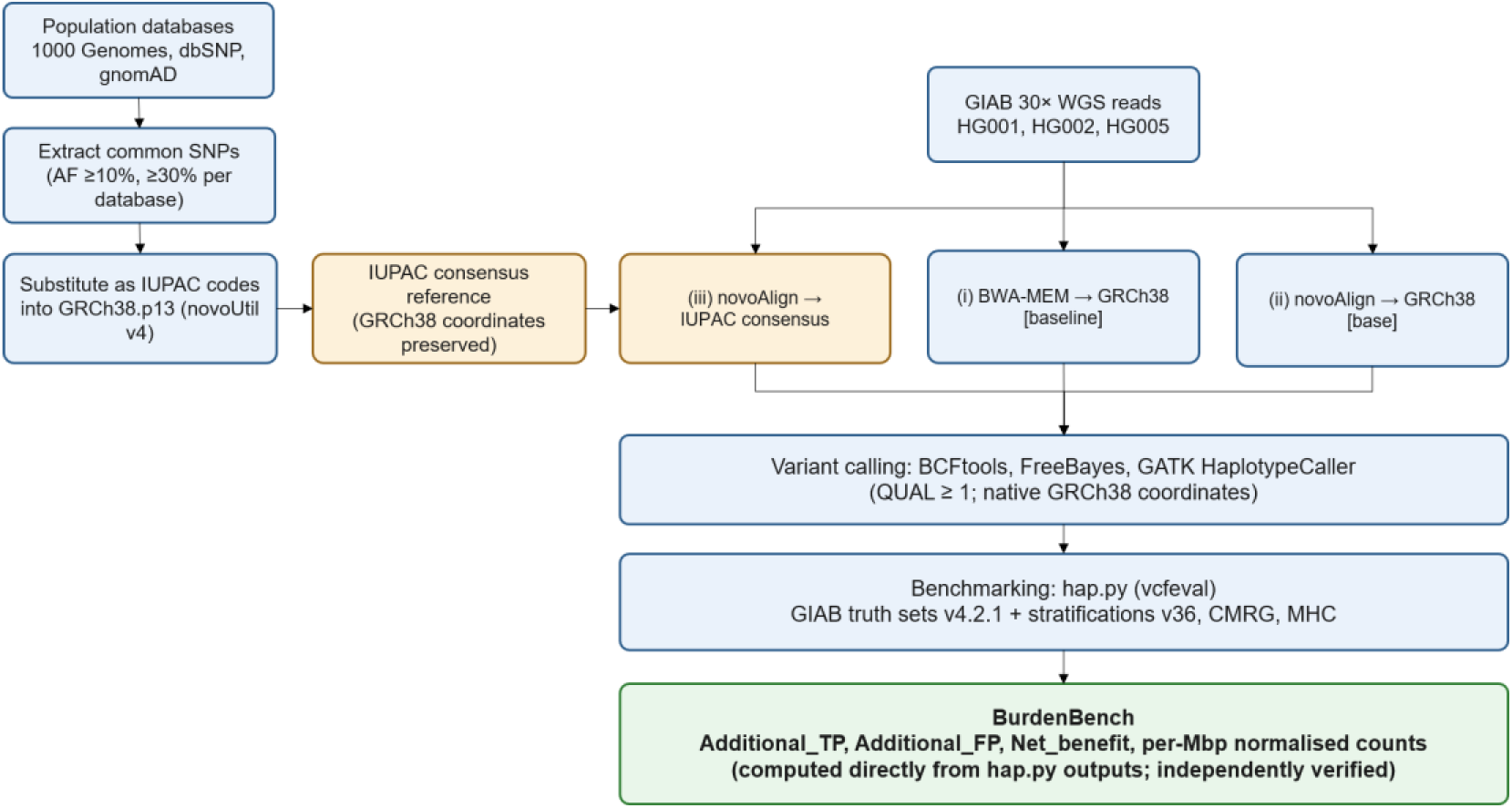
Workflow overview. Common SNPs are extracted from population databases (1000 Genomes, dbSNP, gnomAD) and substituted into GRCh38.p13 using IUPAC degenerate codes via novoUtil. Reads from GIAB samples are aligned using (i) BWA-MEM/GRCh38 (baseline), (ii) novoAlign/GRCh38 (base) or (iii) novoAlign/IUPAC consensus. Variant calling (BCFtools, FreeBayes, GATK HaplotypeCaller) and benchmarking (hap.py, GIAB truth sets) are performed in native GRCh38 coordinates. BurdenBench operates on standard hap.py outputs to compute absolute true-positive and false-positive trade-offs.

### 2.2 Consensus reference generation

We used GRCh38.p13 as the base reference. SNPs were sourced from 1000 Genomes (version 27022019; 1000 Genomes Project Consortium, 2015), dbSNP (build 151) and gnomAD (r3.0). For 1000 Genomes and gnomAD, common variants were defined by allele-frequency thresholds of ≥10% and ≥30% per database; population-specific SNPs (AF ≥10% and ≥30%) were extracted from gnomAD for AFR, ASJ, EAS and NFE (Supplementary Table S5). For dbSNP, variants were filtered using the Common Allele Frequency (CAF) field at two thresholds, CAF ≤90% and CAF ≤70%, approximately corresponding to the ≥10% and ≥30% alternate-allele-frequency thresholds used for the other databases; the dbSNP CAF90 set was also used for the SALT comparison.

Consensus references were constructed by substituting GRCh38 reference alleles with IUPAC degenerate codes using novoUtil (version 4) (Hercus, 2022b). Multi-allelic SNPs were represented using IUPAC codes to preserve allelic diversity while maintaining linear coordinates; consensus references differ from GRCh38 only in base substitution. Variant calling and benchmarking were performed directly in native GRCh38 coordinates without liftover.

### 2.3 Read alignment

PCR-free NovaSeq WGS datasets (∼30× coverage) for GIAB samples HG001 (European, female), HG002 (Ashkenazi Jewish, male) and HG005 (East Asian, male) were obtained from the Google Brain Genomics Sequencing repository (Baid et al., 2020). Reads were mapped to unmodified GRCh38 using BWA-MEM (version 0.7.18) (Li and Durbin, 2009) and novoAlign (version 4) (Hercus, 2022a). For consensus references, alignment used novoAlign, which treats ambiguous reference bases probabilistically, reducing the match penalty for a read base that matches any encoded nucleotide; BWA-MEM does not support IUPAC ambiguous bases, preventing its use with IUPAC consensus references.

### 2.4 Variant calling

SNVs and INDELs were called against GRCh38.p13 using BCFtools (version 1.9) (Danecek and McCarthy, 2017), FreeBayes (version 1.3.1) (Garrison and Marth, 2012) and GATK HaplotypeCaller (version 4) (Van der Auwera et al., 2013), all configured to report variants with QUAL ≥1 without caller-specific filtering, to isolate alignment effects from filtration behaviour. A subset with hard filtering (QUAL ≥30) was separately evaluated to confirm that precision losses at QUAL ≥1 represent conservative upper bounds (Supplementary Table S4).

### 2.5 Benchmarking

Variant call accuracy was quantified with hap.py version 0.3.15 (vcfeval) (Krusche, 2018; Cleary et al., 2015) following established benchmarking guidelines (Krusche et al., 2019), using NIST GIAB truth sets v4.2.1 for HG001, HG002 and HG005 stratified with GIAB stratifications for GRCh38 (v36) (Dwarshuis et al., 2024), the CMRGv1 truth set for HG002 (Wagner et al., 2022b) and the MHC truth set for HG002 (Wagner et al., 2022b). Baseline performance was defined by BWA-MEM alignments to GRCh38. Analyses covered autosomes only; CMRG and MHC annotations are available only for HG002.

### 2.6 BurdenBench implementation

BurdenBench (v1.0.0) computes, from hap.py outputs: Additional_TP = QUERY.TP_consensus − QUERY.TP_baseline; Additional_FP = QUERY.FP_consensus − QUERY.FP_baseline; Net_benefit = Additional_TP − Additional_FP; and per-Mbp counts normalised by confident region size. Derived metrics (F1 gain, TP:FP ratio with Laplace smoothing) are described in the Supplementary Methods. To mitigate potential bias associated with software developed by an author’s employer, primary hap.py outputs and derived burden metrics were independently verified by M.G.R. and L.M.D. from raw TSV files; raw hap.py outputs, intermediate TSVs and the BurdenBench audit trail are available in the project repository.

### 2.7 Comparison to SNP-aware dual-index alignment

We compared IUPAC consensus alignment to SALT (vbeta0.1), a published SNP-aware short-read aligner with a dual-index architecture (CFM-index for the primary reference and RFM-index for alternative SNP alleles; Quan et al., 2021). Both methods used the identical variant set — dbSNP build 151 SNPs with allele frequency ≤90% (CAF90) — so that any performance divergence was attributable to alignment architecture rather than variant database composition or caller choice. SALT was indexed per the authors’ instructions with the standard GRCh38 primary reference and the dbSNP CAF90 variant set; the IUPAC consensus reference for novoAlign was constructed by substituting the same alleles into GRCh38 using novoUtil. SALT alignment used default parameters, outputting SAM format for downstream processing. Variant calling used BCFtools, FreeBayes and GATK HaplotypeCaller against GRCh38.p13 with identical parameters (QUAL ≥1) to the IUPAC consensus condition, and benchmarking used hap.py (vcfeval) with the same GIAB truth sets, stratifications and burden-calculation framework.

### 2.8 Statistical analysis

With only three GIAB samples, all values are descriptive: median [range] for autosomal stratifications (HG001, HG002, HG005) and single-sample values for CMRG and MHC (HG002 only). We report directional consistency (all three samples showing the same sign of effect), percentage-point changes and net benefit. No confidence intervals or P-values are reported; all findings are hypothesis-generating.

## 3 Results

### 3.1 Recall and precision changes by region

Consensus-genome alignment improved variant detection across all three samples, with the largest gains in low-mappability (LM) and segmental-duplication (SD) regions (Table 1; Fig. 2; AS = autosomes, D = difficult regions, LM = low-mappability, SD = segmental duplications, HP = homopolymers, TR = tandem repeats, CMRG = challenging medically relevant genes, MHC = major histocompatibility complex). SNV recall increased by 3.1–3.9 pp and INDEL recall by 4.5–5.8 pp in LM, with gains directionally consistent across all three samples for every caller; precision changes were modestly negative for all callers under the permissive QUAL ≥1 threshold (SNVs: −3.87 to −2.71 pp; INDELs: −3.82 to −2.17 pp), reflecting the absence of post-call filtering. SD showed smaller but consistent SNV gains (1.8–3.1 pp) and INDEL gains (2.4–3.8 pp), with near-neutral INDEL precision. In the MHC (HG002 only), SNV recall rose 2.3–2.9 pp and INDEL recall 3.2–4.4 pp, with GATK showing the largest gains; in CMRG (HG002 only), GATK showed smaller but still positive gains (SNV +0.264 pp, INDEL +0.385 pp). Across all autosomes, SNV recall increased by 0.19–0.24 pp and INDEL recall by 0.1–2.1 pp, with INDEL precision gains of +0.7 to +1.4 pp for BCFtools and FreeBayes.

**Fig. 2.**
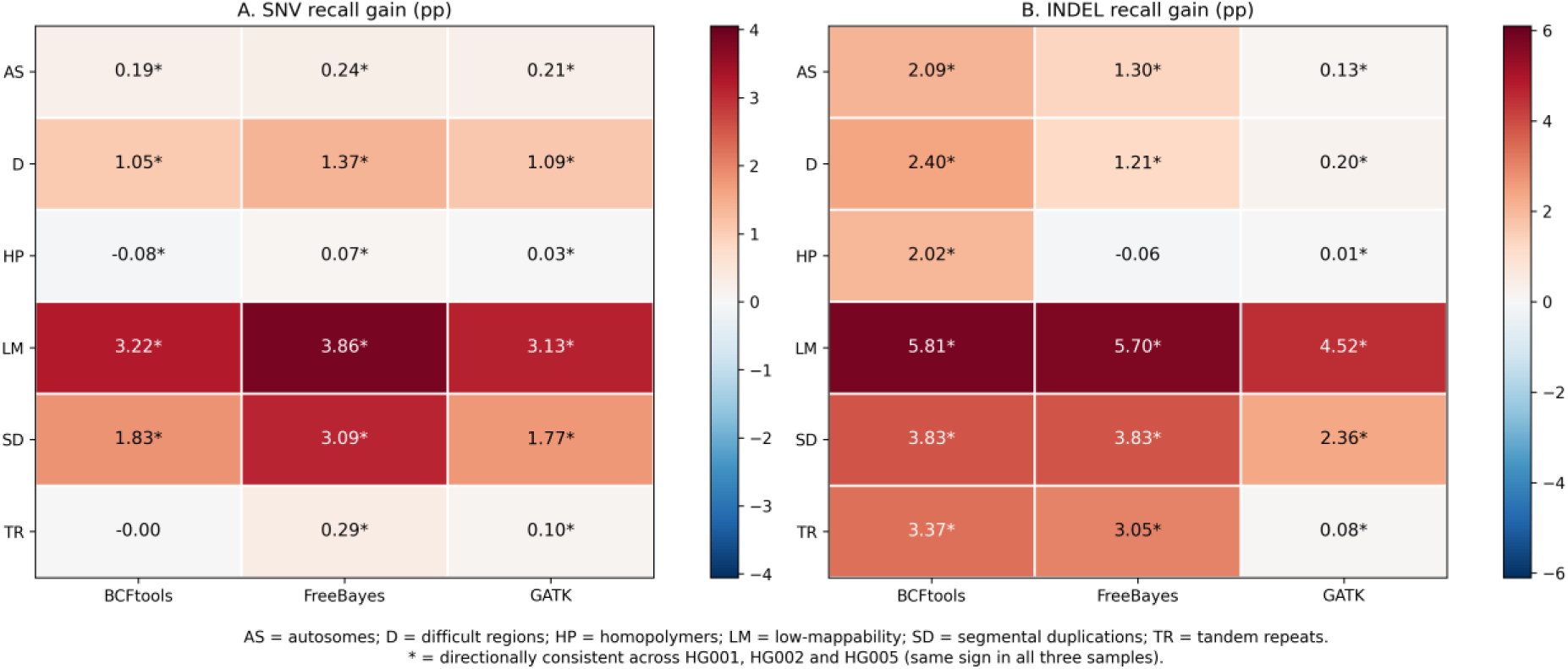
Median recall gain (novoAlign/IUPAC versus BWA-MEM/GRCh38) by GIAB stratification and caller, for (A) SNVs and (B) INDELs. Asterisks denote directional consistency across HG001, HG002 and HG005 (same sign in all three samples).

**Table 1.**
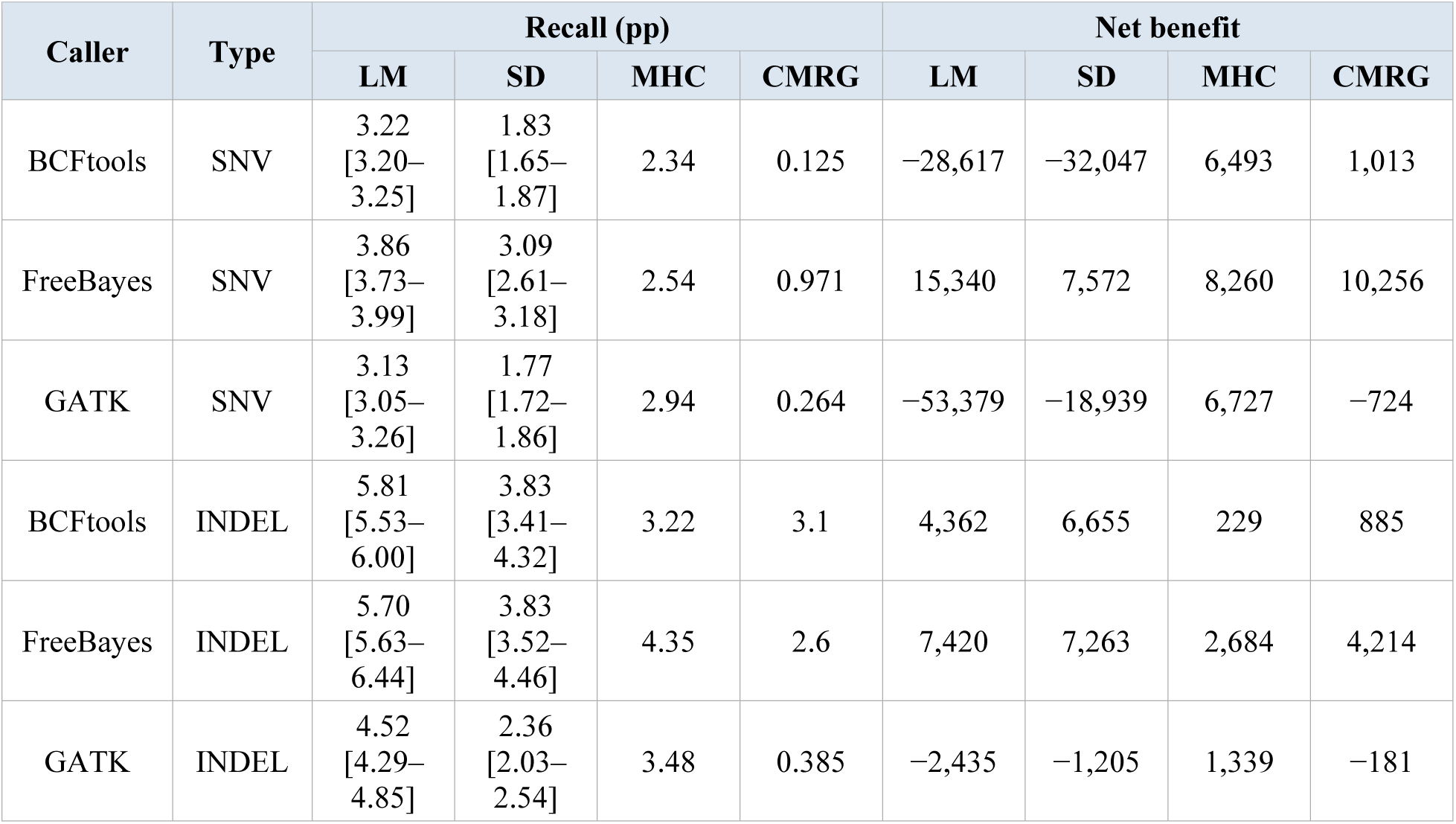
Stratified median recall gain (pp) and net benefit for IUPAC consensus (novoAlign/IUPAC) relative to the BWA-MEM/GRCh38 baseline. Values are median [range] across HG001, HG002 and HG005 for LM and SD; single-sample (HG002) for MHC and CMRG. Net benefit = Additional_TP − Additional_FP. Full results in Supplementary Table S2.

| Caller | Type | Recall (pp) |  |  |  | Net benefit |  |  |  |
| --- | --- | --- | --- | --- | --- | --- | --- | --- | --- |
|  |  | LM | SD | MHC | CMRG | LM | SD | MHC | CMRG |
| BCFtools | SNV | 3.22<br>[3.20–3.25] | 1.83<br>[1.65–1.87] | 2.34 | 0.125 | –28,617 | –32,047 | 6,493 | 1,013 |
| FreeBayes | SNV | 3.86<br>[3.73–3.99] | 3.09<br>[2.61–3.18] | 2.54 | 0.971 | 15,340 | 7,572 | 8,260 | 10,256 |
| GATK | SNV | 3.13<br>[3.05–3.26] | 1.77<br>[1.72–1.86] | 2.94 | 0.264 | –53,379 | –18,939 | 6,727 | –724 |
| BCFtools | INDEL | 5.81<br>[5.53–6.00] | 3.83<br>[3.41–4.32] | 3.22 | 3.1 | 4,362 | 6,655 | 229 | 885 |
| FreeBayes | INDEL | 5.70<br>[5.63–6.44] | 3.83<br>[3.52–4.46] | 4.35 | 2.6 | 7,420 | 7,263 | 2,684 | 4,214 |
| GATK | INDEL | 4.52<br>[4.29–4.85] | 2.36<br>[2.03–2.54] | 3.48 | 0.385 | –2,435 | –1,205 | 1,339 | –181 |

### 3.2 Decomposing aligner and IUPAC contributions

To separate aligner and reference effects, we additionally aligned the same samples to unmodified GRCh38 using novoAlign, creating a base condition (novoAlign/GRCh38) for comparison against both the consensus condition (novoAlign/IUPAC) and the BWA-MEM/GRCh38 baseline. novoAlign alone improved recall over BWA-MEM, particularly for INDELs: in LM regions, SNV recall increased by 0.95–1.6 pp and INDEL recall by 3.5–4.6 pp with the unmodified reference (Fig. 3), indicating that the aligner change itself contributes meaningfully to the overall gain. The IUPAC consensus provided additional recall gains beyond this aligner effect: comparing novoAlign/IUPAC to novoAlign/GRCh38, the additional SNV recall gain from IUPAC encoding was 1.5–2.3 pp in LM and 1.0–1.3 pp for INDELs, with similar patterns in SD, MHC and CMRG. The aligner change therefore drives a substantial portion of INDEL gains, while IUPAC encoding contributes additional SNV-specific improvement. Because BWA-MEM does not support IUPAC ambiguous bases, however, the IUPAC effect could not be isolated independently of aligner choice: IUPAC consensus requires an ambiguity-aware aligner. Precision effects were caller-specific: in LM, BCFtools showed a small additional SNV precision loss (−0.7 pp) with IUPAC beyond the aligner effect, FreeBayes was essentially unchanged (−0.5 pp) and GATK improved (+0.5 pp); for INDELs in LM, BCFtools and GATK improved (+0.7 pp and +0.4 pp) while FreeBayes declined (−0.6 pp).

**Fig. 3.**
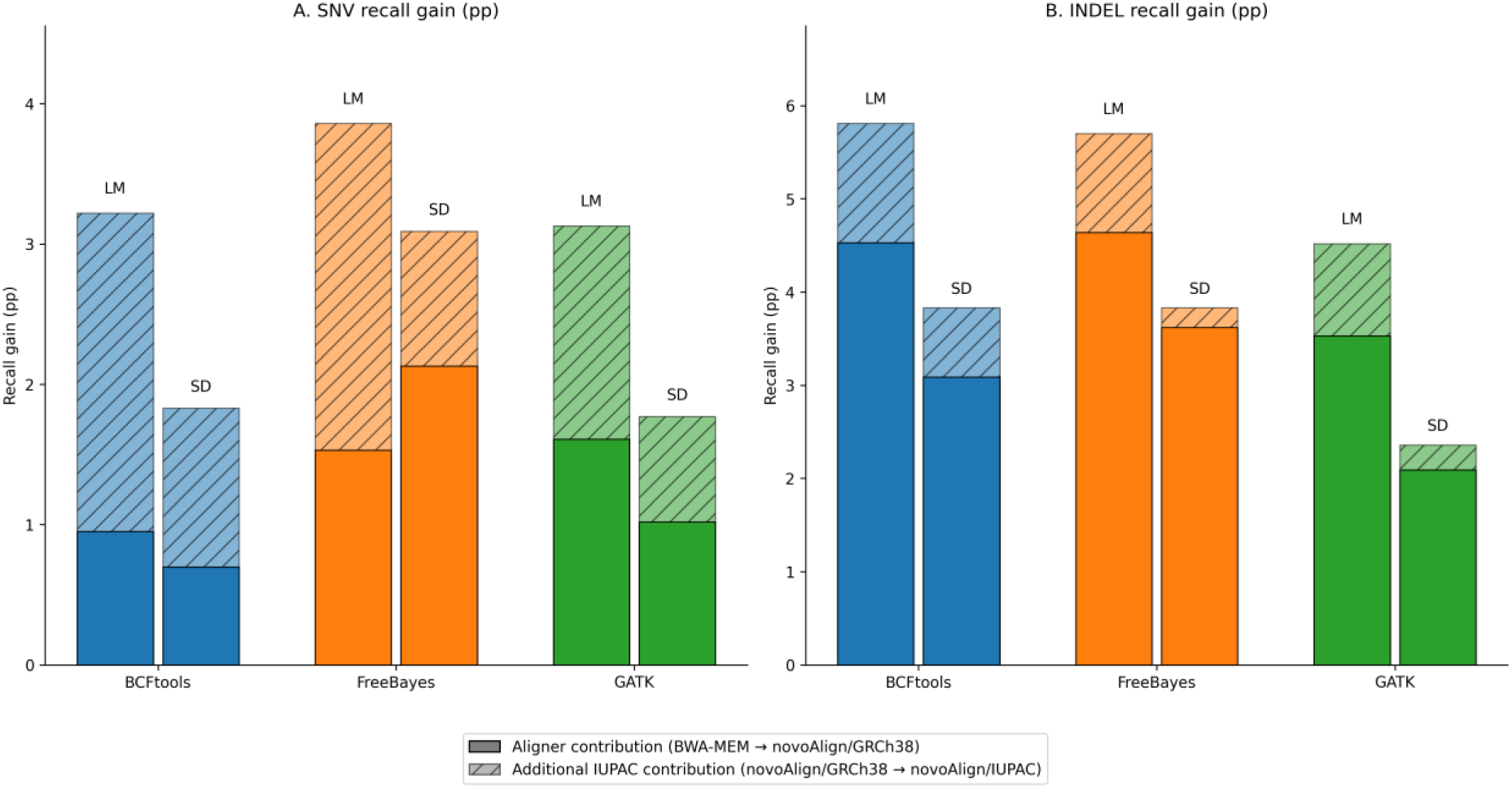
Decomposition of recall gains into aligner contribution (BWA-MEM → novoAlign/GRCh38) and additional IUPAC contribution (novoAlign/GRCh38 → novoAlign/IUPAC), by region (LM, SD) for (A) SNVs and (B) INDELs. Bar segments sum to the total recall gain reported in Table 1. Full decomposition values, including MHC and CMRG, are given in Supplementary Tables S2 and S7; LM values are also reported in the main text (Section 3.2).

### 3.3 BurdenBench: absolute benchmarking and caller-specific trade-offs

Percentage-point recall and precision changes describe relative performance but do not reveal how many variants are gained or lost in absolute terms: a method that improves recall by 3 pp in a 200-Mbp region gains far more true positives than the same improvement in a 2-Mbp region, so per-Mbp normalisation is essential for fair cross-stratification comparison. BurdenBench quantifies these trade-offs by computing, for each caller and stratification, the additional true positives (Additional_TP), additional false positives (Additional_FP), net benefit and per-Mbp-normalised counts directly from hap.py outputs. Applying BurdenBench to the IUPAC consensus results revealed divergent caller-specific profiles invisible to aggregate F1 alone (Table 1; per-sample values in Supplementary Table S1, Fig. S1). In LM, FreeBayes gained 72.5 true positives per Mbp at a cost of 61.6 false positives per Mbp, yielding positive net benefit in all three samples (+15,340); BCFtools and GATK gained comparable numbers of true positives per Mbp (59.6 and 57.3) but incurred more false positives per Mbp (79.1 and 94.1), producing negative net benefit for SNVs (−28,617 and −53,379). For INDELs, net benefit was positive for BCFtools (+4,362) and FreeBayes (+7,420) but negative for GATK (−2,435). In SD, net benefit was strongly positive for FreeBayes (+7,263 INDELs) and moderately positive for BCFtools (+6,655 INDELs), but negative to near-neutral for GATK. In the MHC, GATK achieved positive net benefit for both SNVs (+6,727) and INDELs (+1,339), reflecting fewer false positives per true positive gained than in LM or SD; FreeBayes also showed positive net benefit (SNVs +8,260; INDELs +2,684), while BCFtools showed positive net benefit for SNVs (+6,493) but only modestly for INDELs (+229). In CMRG, INDEL net benefit was positive for BCFtools (+885) and FreeBayes (+4,214), but GATK showed negative net benefit for both variant types, though the absolute numbers of additional variants were small. Across all autosomes, net benefit was strongly positive for BCFtools (+405,175) and FreeBayes (+276,523) on INDELs, but negative for GATK SNVs (−99,379).

### 3.4 Allele-frequency thresholds and population specificity

For 1000 Genomes and gnomAD, raising the allele-frequency threshold from ≥10% to ≥30% decreased SNV recall by 0.01–0.23 pp while improving precision by up to 0.33 pp; for INDELs, BCFtools and FreeBayes showed modest recall increases (0.1–0.5 pp) at the higher threshold, while GATK showed minimal change. For dbSNP, the more selective CAF ≤70% threshold (≈alternate AF ≥30%) generally produced larger recall gains and better net benefit than CAF ≤90% (≈AF ≥10%) for INDELs across all three callers; SNV results were mixed, with BCFtools showing positive net benefit at CAF70 (+6,858) but negative net benefit at CAF90 (−22,685), while FreeBayes and GATK showed larger SNV net benefits at CAF70 than CAF90 in the same direction. These patterns suggest CAF70 may offer a better precision–recall balance for routine use (full results in Supplementary Table S3; Fig. S2). A single pan-human consensus performed similarly to population-specific references (AFR, ASJ, EAS, NFE), with differences ≤0.2 pp across all gnomAD stratifications.

### 3.5 Comparison to SNP-aware dual-index alignment

SALT produced consistent and severe performance degradation relative to the BWA-MEM/GRCh38 baseline across all three callers and all stratifications (Table 2; Supplementary Table S6). For SNVs, all callers showed negative recall across every stratification (BCFtools −18.3 to −26.4 pp; FreeBayes −32.0 to −43.2 pp; GATK −5.4 to −9.2 pp), with corresponding precision losses of −9.6 to −61.8, −17.4 to −58.9 and −22.4 to −69.7 pp, respectively; SNV net benefit was negative for all callers in every stratification without exception. The most extreme losses occurred in low-mappability regions (GATK: SNV recall −8.46 pp, precision −69.7 pp, net benefit −624,199). For INDELs, BCFtools showed apparent recall gains in some strata (e.g. tandem repeats, +8.4 pp) but with severe precision losses (−1.6 to −29.2 pp), giving negative net benefit in five of six stratifications; FreeBayes and GATK showed INDEL recall losses across all stratifications with consistently negative net benefit. In contrast, IUPAC consensus alignment with novoAlign achieved modest recall gains (+0.2 to +3.5 pp) with precision changes ranging from −0.1 to −9.1 pp across the same stratifications, and net benefit substantially less negative than SALT for both SNVs (−67,393 versus −479,808 to −624,199 for SALT in the worst strata) and INDELs (−2,598). All three callers underperformed even the unadjusted BWA-MEM/GRCh38 baseline with SALT, confirming that its performance deficit is architectural rather than caller-specific.

**Table 2.** SNV recall and precision gains (pp) for novoAlign/IUPAC and SALT (vbeta0.1) relative to the BWA-MEM/GRCh38 baseline. Both methods used the dbSNP build 151 CAF90 variant set. INDEL results in Supplementary Table S6.

| Caller | Type | Region | Recall gain (pp) |  | Precision gain (pp) |  |
| --- | --- | --- | --- | --- | --- | --- |
|  |  |  | novoAlign/IUPAC | SALT | novoAlign/IUPAC | SALT |
| BCFtools | SNV | AS | 0.49 | −21.7 | −0.52 | −12.6 |
| BCFtools | SNV | LM | 3.22 | −26.4 | −3.87 | −61.8 |
| BCFtools | SNV | SD | 1.83 | −18.3 | −3 | −9.6 |
| FreeBayes | SNV | AS | 0.63 | −43.2 | 0.74 | −17.4 |
| FreeBayes | SNV | LM | 3.86 | −32 | −2.71 | −58.9 |
| FreeBayes | SNV | SD | 3.09 | −38.6 | −1.41 | −38.2 |
| GATK | SNV | AS | 0.49 | −9.19 | −0.87 | −29.1 |
| GATK | SNV | LM | 3.13 | −8.46 | −4 | −69.7 |
| GATK | SNV | SD | 1.77 | −5.42 | −2.03 | −60.7 |

## 4 Discussion

### 4.1 Principal findings and mechanism

Aligning short-read data to an IUPAC consensus reference, combined with an ambiguity-aware aligner, improved recall in the regions where clinical pipelines struggle most: low-mappability regions, segmental duplications, the MHC and medically relevant genes. SNV recall rose by 3.1–3.9 pp and INDEL recall by 4.5–5.8 pp in the most challenging areas, with gains directionally consistent across three individuals and three callers for the autosomal stratifications, indicating that the effect is not specific to a single pipeline or genetic background. The improvement arises from reduced mismatch penalties at common variant sites: aligners that support IUPAC codes score ambiguous positions as matches, raising mapping quality in polymorphic regions (Kaminow et al., 2022), and the decomposition analysis shows that the aligner change (BWA-MEM to novoAlign) drives a substantial portion of INDEL gains while IUPAC encoding contributes additional SNV-specific improvement. These are descriptive findings from three samples for autosomal stratifications and from a single sample (HG002) for CMRG and MHC; they identify a pattern warranting validation and do not establish clinical efficacy or population-level generalisability.

### 4.2 BurdenBench as a reusable framework

The central methodological contribution of this work is BurdenBench, a reproducible framework for absolute true-positive and false-positive quantification. Traditional benchmarking relies on precision, recall and F1, which summarise performance but obscure the absolute counts that determine laboratory workload and interpretation cost; BurdenBench addresses this by reporting Additional_TP, Additional_FP, net benefit and per-Mbp-normalised counts directly from hap.py outputs, requiring no modification to hap.py or truth sets. It can therefore be applied retrospectively to published benchmarking data and prospectively to new methods, including evaluations of alignment strategies, caller parameters and filtering thresholds. The per-Mbp normalisation proved essential here: without it, comparing absolute counts across regions of vastly different size would be misleading, and the hypothesis that consensus references may be most cost-effective when applied selectively to difficult regions would remain untestable.

### 4.3 Caller-specific trade-offs and clinical priorities

The divergent burden profiles revealed by BurdenBench carry practical implications. FreeBayes achieved the most favourable balance between additional true positives and false positives across most regions; BCFtools showed strong INDEL net benefit but higher SNV false-positive rates; and GATK generated more false positives per true positive gained in LM and SD, but achieved positive net benefit in the MHC and showed consistent recall gains across all clinically relevant stratifications. This suggests that selective application of IUPAC consensus to these regions, paired with existing VQSR or hard-filtering pipelines, is a practical deployment option for GATK users. The net-benefit metric, which weights true and false positives equally, does not capture an important clinical asymmetry: a missed pathogenic variant may remain undetected until persistent symptoms or failed diagnosis triggers reanalysis or resequencing, incurring diagnostic delay and cost, whereas a false positive can be flagged by manual review, Sanger confirmation or segregation analysis. Laboratories may therefore reasonably accept higher false-positive rates to maximise sensitivity in challenging regions, particularly for INDELs where recall gains were largest. The MHC results illustrate this prioritisation most clearly: GATK achieved approximately six true variants detected for every false call introduced, and existing VQSR or hard-filtering pipelines can address the precision cost. GATK’s CMRG net benefit, by contrast, was negative for both variant types with default parameters, suggesting that selective application or parameter tuning may be warranted for this region specifically.

### 4.4 Tunable trade-offs via allele-frequency thresholds

Across all three databases, the lower AF threshold (≥10% for 1000 Genomes and gnomAD; CAF ≤90% for dbSNP) maximises SNV recall in difficult regions and is preferable for discovery-oriented pipelines, while the higher threshold (≥30% or CAF ≤70%) offers a modest precision gain at small recall cost and may suit confirmatory workflows. The dbSNP CAF ≤70% threshold showed a particularly favourable trade-off profile, with BCFtools SNV net benefit positive at CAF70 but negative at CAF90, and INDEL net benefit consistently larger at CAF70 across all callers. Because we deliberately disabled caller-specific filters (QUAL ≥1) to isolate alignment effects, the precision losses reported here are conservative upper bounds. Standard hard filtering (QUAL ≥30) improved precision for most caller–variant combinations in our exploratory analysis, with BCFtools SNV net benefit turning positive (Supplementary Table S4); routine application of calibrated filters or machine-learning post-filters in clinical pipelines is expected to recover precision with limited recall impact.

### 4.5 Global versus population-specific consensus

Pan-human and population-specific consensuses performed within 0.2 pp across all callers and thresholds, extending a similar finding from RNA-seq (Kaminow et al., 2022). With only three samples, population-specific differences are indistinguishable from sampling variation, and a single global consensus remains a practical starting point pending larger truth sets.

### 4.6 Clinical relevance and validation needs

Many clinically important genes reside in low-mappability and segmental-duplication regions where GRCh38 representation leads to misalignment or missed calls, a problem also documented using orthogonal linked- and long-read validation (Wagner et al., 2022a). We observed SNV recall gains in CMRG of up to 1.0 pp (648 additional true positives for the GATK IUPAC median) and INDEL recall gains of up to 3.1 pp (3,189 additional true positives for BCFtools). In the MHC, where variability is high and clinical utility spans autoimmune disease association and cancer immunogenomics, SNV recall gains were 2.3–2.9 pp (6,471–8,137 additional true positives) and INDEL recall gains 3.2–4.4 pp (1,618–2,186 additional true positives), consistent with bias reduction when the reference carries common alleles rather than a single haplotype (Lin et al., 2024). Reference bias may disproportionately affect individuals whose haplotypes diverge most from GRCh38, but the clinical impact remains poorly quantified (Ballouz et al., 2019). Our three samples cover European, Ashkenazi Jewish and East Asian ancestries, suggesting the pan-human consensus may be broadly compatible; direct validation on additional ancestries, including African populations, is needed once GIAB truth sets with the required stratifications become available beyond HG001, HG002 and HG005.

### 4.7 Architectural lessons: IUPAC versus dual-index alignment

Our controlled comparison to SALT (Quan et al., 2021), using the identical SNP set, reference genome and all three variant callers, indicates that among linear-reference variant-aware alignment strategies, the mechanism of variant incorporation can determine performance. Both IUPAC consensus and SALT incorporate common-variant information into a linear reference but differ architecturally: IUPAC encoding modifies the reference sequence in place using degenerate bases, whereas SALT builds a dual index. SALT’s design produced severe precision losses (−9.6 to −69.7 pp) and recall losses (−5.4 to −43.2 pp) across callers and stratifications, with all three callers underperforming even the variant-naive BWA-MEM baseline, confirming that the deficit is architectural rather than specific to any one caller’s haplotype model. We acknowledge limitations of this comparison: SALT was originally designed for alignment accuracy rather than GIAB benchmarking, its evaluation metrics (perfect alignment rate) differ from the precision–recall framework used here, and its performance with its own recommended variant-calling pipeline was not evaluated. The broader aligner landscape reinforces our conclusion: DRAGEN’s graph mapper uses IUPAC codes for isolated SNPs but constructs population alternate contigs for haplotypes at index-building time (Illumina, 2021), making its IUPAC support architecturally distinct from novoAlign’s scoring-based approach, and mrFAST’s SNP-aware mode produced a segmentation fault on our datasets, a known unresolved issue (sfu-compbio/mrsfast issue #14). The available evidence consistently favours approaches that preserve linear reference structure with probabilistic variant scoring.

### 4.8 Positioning within the broader reference landscape

IUPAC consensus alignment is not a replacement for T2T-CHM13 or HPRC pangenome references, which resolve regions inaccessible to GRCh38 (Nurk et al., 2022; Liao et al., 2023); its practical advantage is operational simplicity: it preserves GRCh38 linear coordinates and GA4GH benchmarking compatibility without requiring pipeline re-validation. Direct comparison to T2T/HPRC references is needed to define where IUPAC consensus offers independent value.

### 4.9 Limitations

**1.** Sample size. Only three GIAB samples were analysed for autosomal stratifications. This reflects availability rather than experimental design: GIAB truth sets with the required stratifications (v36) are currently limited to HG001, HG002 and HG005 for autosomal regions, and CMRG and MHC annotations are available only for HG002. All reported autosomal values are descriptive medians and ranges; CMRG and MHC results are single-sample values. No confidence intervals, P-values or standard errors are reported. The directional consistency across the three samples supports the robustness of the observed pattern, but it remains a pattern, not an established effect.
**2.** Aligner–reference confounding. The IUPAC effect cannot be fully decoupled from the aligner change, because BWA-MEM does not support IUPAC ambiguous bases. The decomposition analysis (novoAlign/GRCh38 versus novoAlign/IUPAC) partially addresses this, and the SALT comparison suggests that preserving linear reference structure is architecturally consequential; however, novoAlign remains the only publicly documented aligner supporting true IUPAC degenerate base scoring at the reference-index level, and validating the IUPAC encoding effect with a fully independent aligner remains an open experimental goal pending software maintenance or new tool development.
**3.** GATK parameter optimisation. GATK showed higher false-positive rates than BCFtools or FreeBayes in most stratifications, though recall gains were consistent across all regions and net benefit was positive in the MHC. We did not investigate parameter optimisation that might improve the precision–recall trade-off; laboratories using GATK should evaluate IUPAC consensus with their existing filter chains before drawing conclusions about suitability.
**4.** No pangenome comparison. We did not benchmark against T2T-CHM13 or HPRC pangenome references; claims that IUPAC consensus is a pragmatic bridge are speculative without direct comparison.
**5.** No sex-chromosome analysis; performance on X-linked genes (e.g. DMD, F8) is unknown.
**6.** Permissive calling (QUAL ≥1); precision losses are conservative upper bounds.
**7.** IUPAC substitution does not correct GRCh38 assembly errors.

## 5 Availability and implementation

BurdenBench (version 1.0.0) is available at https://github.com/akzam/BurdenBench under the MIT licence. It requires Python 3.7 or higher, pandas and numpy, and can be installed via pip or conda. Raw hap.py benchmarking outputs are provided in the repository to allow independent recomputation without requiring a novoAlign licence. Sequencing data for GIAB samples HG001, HG002 and HG005 are available from the Google Brain Genomics Sequencing repository (Baid et al., 2020; doi: 10.1101/2020.12.11.422022). GIAB truth sets (v4.2.1) and stratifications are available from NIST (https://www.nist.gov/programs-projects/genome-bottle; Dwarshuis et al., 2024). CMRG and MHC truth sets are available from Wagner et al. (2022b). Population variation resources used were the 1000 Genomes Project, dbSNP (build 151; Sherry et al., 2001) and gnomAD v3 (Karczewski et al., 2020). novoAlign version 4 and novoUtil version 4 are commercial software (Novocraft Technologies; Hercus, 2022a, 2022b); academic trial licences are available at no cost.

## Funding

This study was supported by National and Health Medical Research Council of Australia (Senior Research Fellowship: 1104718; Project Grant: 1125523 to L.M.D.) and institutional funding from the University of South Australia (now Adelaide University). A.S. was supported by an IRTS fee-waiver scholarship from the University of South Australia (now Adelaide University) and is an employee of Novocraft Technologies.

## Conflict of interest statement

A.S. is a PhD candidate at Adelaide University and an employee of Novocraft Technologies, the developer of novoAlign and novoUtil, and receives salary from Novocraft. This study was designed and conducted as part of A.S.’s doctoral research under the independent academic supervision of M.G.R. and L.M.D. Novocraft Technologies provided no funding, had no role in study design, data collection, analysis, interpretation or manuscript preparation, and exercised no editorial control. M.G.R. and L.M.D. had full access to all raw benchmarking outputs, independently verified the primary burden calculations, and retain final authority over manuscript content. All other authors declare no competing interests.

## Data availability

All data underlying this article are publicly available as described in Section 5 (Availability and implementation) and are cited in the text. No new sequencing data were generated.

## Acknowledgements

The authors thank Adelaide University for laboratory funding and the tuition scholarship for A.S., and Novocraft Technologies for providing free access to novoAlign and novoUtil and for clarifications regarding the IUPAC scoring scheme. Manuscript editing assistance was provided using Claude (Anthropic) as an AI writing aid; all scientific content, data analysis and conclusions are the sole responsibility of the authors.

## Author contributions

Conceptualisation, A.S.; Methodology, A.S.; Software, A.S.; Formal analysis, A.S.; Investigation, A.S.; Resources, M.G.R. and L.M.D.; Data curation, A.S.; Writing — original draft, A.S.; Writing — review and editing, M.G.R. and L.M.D.; Visualisation, A.S.; Supervision, M.G.R. and L.M.D.; Funding acquisition, M.G.R. and L.M.D.

## Notes

https://github.com/akzam/BurdenBench

