## Supplemental Figures for "IUPAC Consensus References Improve Short-Read Variant Detection in Clinically Challenging Regions: A Stratified Benchmarking Study with BurdenBench"

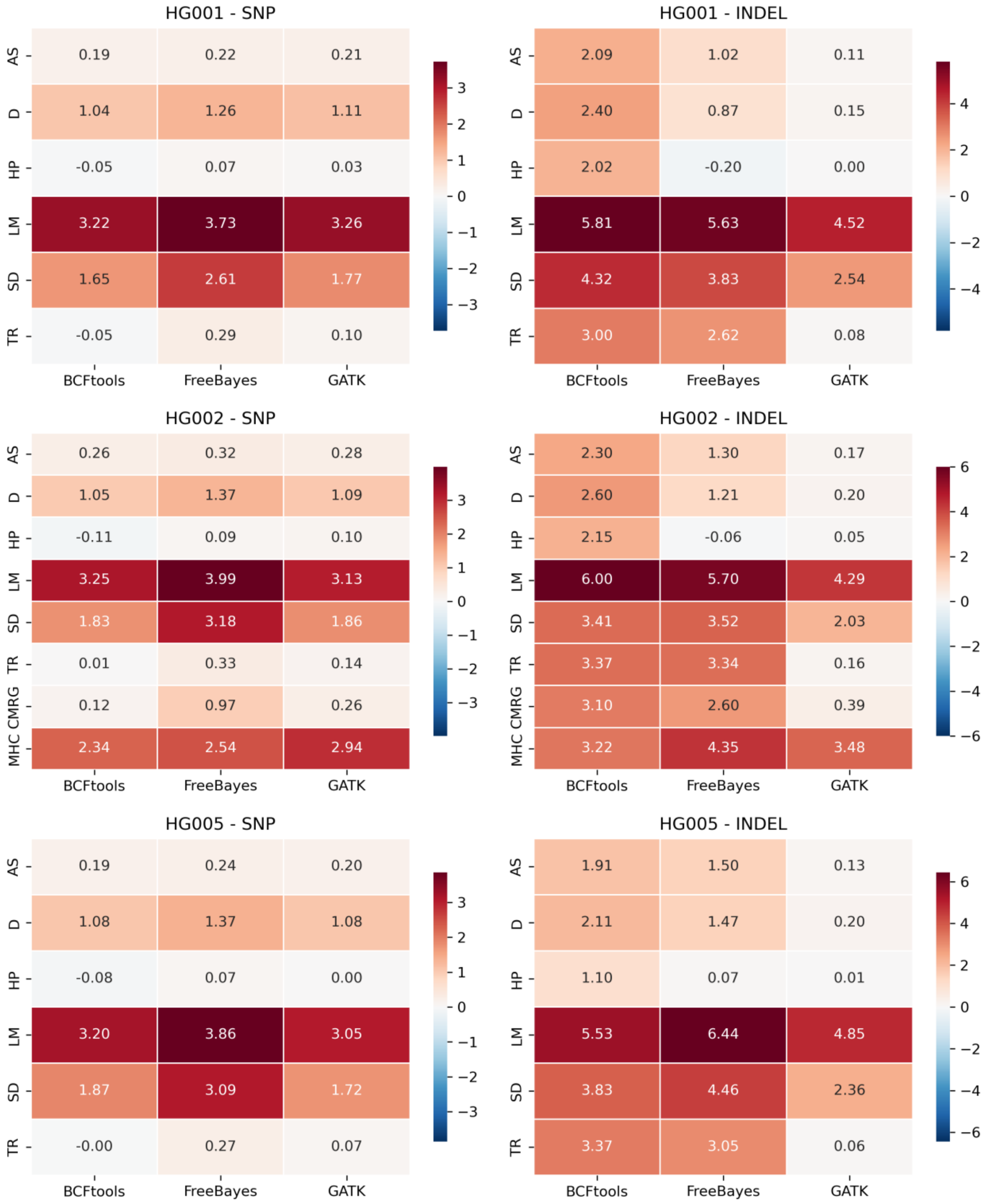


Fig. S1. Heatmap of recall gain (novoAlign/IUPAC versus BWA-MEM/GRCh38) by sample, GIAB stratification, caller and variant type. See main text Section 3.1 for stratification abbreviations.

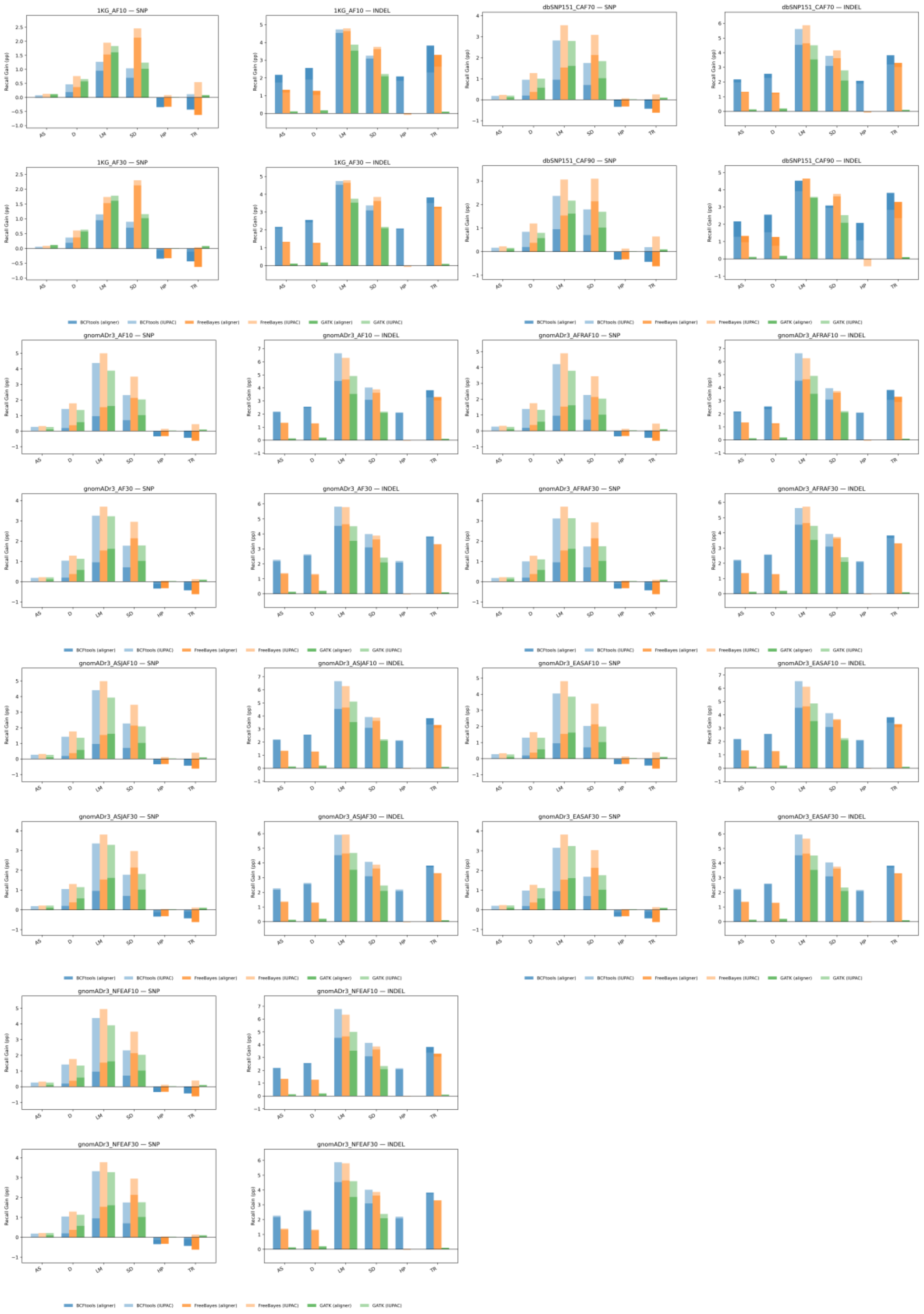


Fig. S2. Stacked bar chart showing total recall gain decomposed into aligner contribution (BWA-MEM → novoAlign/GRCh38) and IUPAC contribution (novoAlign/GRCh38 → novoAlign/IUPAC) across population-frequency conditions and GIAB stratifications (AS, D, LM, SD, HP, TR).
