## Supplemental Methods for "IUPAC Consensus References Improve Short-Read Variant Detection in Clinically Challenging Regions: A Stratified Benchmarking Study with BurdenBench"

### Complete mathematical derivation of burden framework formulas

#### 1. Percentage-point derivation

All percentage-point (pp) changes reported in the main text and tables are derived directly from hap.py METRIC.Recall and METRIC.Precision values, which are output as proportions on a 0-1 scale:

Recall_gain_pp = (METRIC.Recall_consensus - METRIC.Recall_baseline) * 100

Precision_gain_pp = (METRIC.Precision_consensus - METRIC.Precision_baseline) * 100

Raw 0-1 values are preserved in the output (Recall_base_raw, Recall_cons_raw, Precision_base_raw, Precision_cons_raw) so that every pp value can be independently recalculated. A machine-precision drift check verifies that gain_pp equals (cons_raw - base_raw) * 100 (tolerance > 1e-10); any non-zero drift triggers a validation warning.

#### 2. Absolute burden metrics

For each consensus condition relative to the BWA-MEM/GRCh38 baseline, the framework computes the following from hap.py count columns:

Additional_TP = QUERY.TP_consensus - QUERY.TP_baseline

Additional_FP = QUERY.FP_consensus - QUERY.FP_baseline

Additional_FN = (TRUTH.TOTAL_consensus - QUERY.TP_consensus) - (TRUTH.TOTAL_baseline - QUERY.TP_baseline)

Net_benefit = Additional_TP - Additional_FP

Interpretation: Net benefit is a signed arithmetic summary of the detection trade-off: positive values indicate more true variants detected than false calls introduced; negative values indicate the reverse. This is a descriptive count summary, not a weighted assessment of clinical utility.

TRUTH.TOTAL mismatch warning: If TRUTH.TOTAL differs between baseline and consensus hap.py outputs (e.g., due to different confident region definitions), Additional_FN becomes unreliable because the change in false negatives is confounded by a change in truth-set size. The framework logs a warning when such mismatches are detected.

#### 3. TP:FP ratio (heuristic, for reference only)

The framework computes two ratio variants:

Raw ratio:

TP_FP_ratio_raw = Additional_TP / Additional_FP

This is undefined (NaN) when both numerator and denominator are zero, positive infinity when FP = 0 and TP > 0, and negative infinity when FP = 0 and TP < 0.

Laplace-adjusted ratio:

To avoid division-by-zero and infinite values while preserving rank order, a conditional pseudocount adjustment is applied:

If Additional_TP = 0 AND Additional_FP = 0: NaN (no meaningful ratio)

If Additional_FP >= 0: (Additional_TP + 1) / (Additional_FP + 1)

If Additional_FP < 0 (i.e., fewer false positives in consensus):

Additional_TP / Additional_FP (raw ratio, preserving negative sign)

Caveats: Laplace (+1) smoothing biases the ratio toward 1.0, especially for small counts (bias can exceed 30% when per-Mbp counts < 2). The adjusted ratio is therefore a heuristic summary, not a formal statistical estimator. Primary interpretation should rely on the raw Additional_TP and Additional_FP counts. A formatted display string maps edge cases for readability (e.g., "inf (pure gain)", "0 (cost only)").

**4. Region-size normalisation**

To enable comparison across stratifications of different genomic extents, counts are normalised by the baseline confident region size:

Additional_TP_per_Mbp = Additional_TP / (Subset.IS_CONF.Size_baseline / 1,000,000)

Additional_FP_per_Mbp = Additional_FP / (Subset.IS_CONF.Size_baseline / 1,000,000)

The baseline region size is used for both metrics to ensure the delta is referenced to a common denominator. If the consensus Subset.IS_CONF.Size differs from the baseline by >5% (relative), the framework emits a warning, because per-Mbp normalisation assumes comparable confident regions.

Per-100-truth variants:

As an alternative normalisation, the framework also reports:

$$Additional\_TP\_per\_100\_truth = (Additional\_TP / TRUTH.TOTAL\_baseline) * 100$$

$$Additional\_FP\_per\_100\_truth = (Additional\_FP / TRUTH.TOTAL\_baseline) * 100$$

**5. F1-score computation**

F1 is computed as the harmonic mean of recall and precision:

$$F1 = (2 * Recall * Precision) / (Recall + Precision)$$

with protection against division by zero (F1 = NaN when Recall + Precision = 0).

The percentage-point change is:

$$F1\_gain\_pp = (F1\_consensus - F1\_baseline) * 100$$

**6. Aggregation rules**

The framework supports three aggregation modes, all of which exclude Sample from the grouping keys because samples are the unit of replication.

**6.1. Per-sample computation**

For each sample, burden metrics are computed directly from the pairwise merge of baseline and consensus hap.py outputs. No averaging occurs within a sample; every stratification row (Type x Subtype x Subset x Filter x Genotype) retains its independent value.

**6.2. Cross-sample median and range**

Across the three GIAB samples (HG001, HG002, HG005), the framework reports:

- Median: the median of per-sample values for each metric.
- Range: [minimum - maximum] of per-sample values.
- Count: number of samples contributing to the group.

Grouping columns default to Caller, Condition, Type, Subset, Filter, Genotype.

This produces the stratified summary tables (e.g., Supplemental Table S2).

**6.3. Overall metrics (summed counts, not averaged)**

For "overall" summaries that collapse across stratification rows (e.g., all Filter/Genotype/Subtype values combined), the framework SUMS counts within each sample first, then recomputes recall, precision, and F1 from the aggregated counts:

Recall_overall  = sum(QUERY.TP) / sum(TRUTH.TOTAL)

Precision_overall = sum(QUERY.TP) / sum(QUERY.TP + QUERY.FP)

Averaging recall or precision across stratification rows is mathematically incorrect when truth-set sizes differ, so the framework strictly recomputes from summed counts. Per-Mbp and per-100-truth metrics in overall outputs are most reliable when aggregating non-overlapping strata.

The Condition value 'All-Conditions', reported in Supplemental Table S2 and used for the headline stratified results in the main text and Table 1, applies this overall-metrics rule (Section 6.3) across the population-frequency/database dimension rather than the Filter/Genotype/Subtype dimension: per-sample TP and FP counts are summed across all 14 named population-frequency conditions (Supplemental Table S3: 1KG AF10/30, dbSNP CAF70/90, and gnomAD pan-human and population-specific AF10/30), then recall, precision and net benefit are recomputed from the summed counts before the cross-sample median (Section 6.2) is taken. It does not include the novoAlign/GRCh38 aligner-only 'base' condition (Supplemental Table S7), which is reported separately.

#### 6.4. Median-of-medians (sensitivity analysis)

As a robustness check, the framework can compute the median across all stratification rows WITHIN each sample, then report the median [min-max] of those per-sample medians.

This gives equal weight to each sample regardless of how many stratification rows it contains, and can be compared with the direct median (Section 6.2) to assess aggregation-method sensitivity.

#### 6.5. Duplicate handling

If the manifest contains multiple files for the same sample + caller + condition combination (e.g., technical replicates), the framework averages them by median across duplicates before cross-sample aggregation.

### 7. Validation checks

Each analysis run performs the following checks. Failures are logged; fatal errors halt execution.

1. File access: Baseline and consensus files must exist, be readable, and non-empty.
2. Column completeness: hap.py outputs must contain: Type, Subtype, Subset, Filter, Genotype, METRIC.Recall, METRIC.Precision, QUERY.FP, QUERY.TP, TRUTH.TOTAL, Subset.IS_CONF.Size.
3. Merge-key integrity: No missing values in stratification keys (Type, Subtype, Subset, Filter, Genotype), ensuring unambiguous baseline-consensus pairing.

4. Numeric type correctness: All quantitative columns must be numeric (not string or object dtypes).
5. Value sanity: No negative QUERY.FP or QUERY.TP counts. Recall and precision values outside [0, 1] trigger warnings (they are not fatal, because hap.py may emit boundary values in edge cases).
6. Drift check: Recall_gain_pp and Precision_gain_pp are cross-checked against (cons_raw - base_raw) * 100. Tolerance: 1e-10.
7. Cross-check (FN conservation): The framework warns if TRUTH.TOTAL differs between baseline and consensus, because in that case the expected identity Additional_TP + Additional_FN = TRUTH.TOTAL_consensus - TRUTH.TOTAL_baseline is violated.
8. Region-size drift: If Subset.IS_CONF.Size differs by >5% between baseline and consensus, a warning is emitted because per-Mbp normalisation assumes comparable denominators.

**8. Reproducibility audit trail**

Every batch run generates a metadata file ({prefix}_reproducibility.txt) documenting:

1. Software version: run_BurdenBench.py v1
2. Input provenance: Manifest file path, list of baseline and consensus hap.py files, and per-file checksums (if available).
3. Formula versions: Exact formulas used for pp derivation, burden metrics, Laplace smoothing, F1 computation, and normalisation.
4. Validation results: Which checks were performed and whether they passed or produced warnings.
5. Coverage matrix: A table of Sample x Caller x Condition coverage indicating which comparisons were successfully processed, which failed, and which were averaged from duplicates.
6. Aggregation parameters: Grouping columns, overall levels, and median-of-medians levels requested.
